# The Endoplasmic Reticulum pool of Bcl-xL dampens the Unfolded Protein Response through IP3R-dependent Calcium Release

**DOI:** 10.1101/2021.01.27.428229

**Authors:** Lea Jabbour, Trang Nguyen, Rudy Gadet, Olivier Lohez, Ivan Mikaelian, Philippe Gonzalo, Thomas Luyten, Mounira Chalabi-Dcha, Geert Bultynck, Ruth Rimokh, Germain Gillet, Nikolay Popgeorgiev

**Author notes:** equal contribution (Trang Nguyen, Rudy Gadet). these authors share senior authorship (Ruth Rimokh, Germain Gillet, Nikolay Popgeorgiev).

## Abstract

Apoptosis plays a role in cell homeostasis in both normal development and disease. Bcl-xL, a member of the Bcl-2 family of proteins, regulates the intrinsic mitochondrial pathway of apoptosis. It is overexpressed in several cancers. Bcl-xL has a dual subcellular localization and is found at the mitochondria as well as the endoplasmic reticulum (ER). However, the biological significance of its ER localization is unclear. In order to decipher the functional contributions of the mitochondrial and reticular pools of Bcl-xL, we generated genetically modified mice expressing exclusively Bcl-xL at the ER, referred to as ER-xL, or the mitochondria, referred to as Mt-xL. By performing cell death assays, we showed that ER-xL MEFs show increased vulnerability to apoptotic stimuli but are more resistant to ER stress. Furthermore, ER-xL MEFs demonstrated a reduced expression of the Unfolded Protein Response (UPR) markers upon ER stress and displayed reduced inositol trisphosphate receptor (IP3R)-mediated ER calcium release. Collectively, our data show that upon ER stress, Bcl-xL negatively regulates IP3R-mediated calcium flux from the ER, which prevents ER calcium depletion and maintains the UPR and subsequent cell death in check. This work reveals a moonlighting function of Bcl-xL at the ER, apart from its cliché regulation of apoptosis.

## Introduction

Bcl-xL is a member of the famous Bcl-2 family of proteins ^1^. This family controls the fate of cells by regulating apoptosis. It is composed of numerous proteins that are classified, based on their structure and function. Multidomain Bcl-2 homologs, which comprise cell death inhibitors, such as Bcl-2 and Bcl-XL, and accelerators, such as Bax, possess up to four homology domains referred to as Bcl-2 homology (BH1-4) domains. BH3-only proteins (Bad, Bid, Puma…), which only regroup death accelerators, are also considered as Bcl-2 homologs, even though being from different phylogenic origins^2^. Proapoptotic proteins induce mitochondrial outer membrane permeabilization (MOMP), a key checkpoint upstream of the activation of the caspase cascade resulting in apoptosis ^3^.

Dysregulations affecting the network of Bcl-2 homologs might have detrimental outcomes such as auto-immune or cancer diseases. Indeed, Bcl-xL is upregulated in hepatocellular and renal carcinoma ^4,5^ as well as pancreatic cancer ^6^. Furthermore, Bcl-xL was shown to play a role in the invasion of human malignant glioma ^7^ and colorectal cancer cells ^8^. However, in breast cancer, there is evidence that Bcl-xL does not affect tumor growth but increases metastatic potential, independent from apoptosis ^9,10^.

Throughout the years, non-canonical functions of the Bcl-2 family of proteins have been progressively unmasked. Indeed, besides their mitochondrial localization, most of the Bcl-2 proteins translocate to other subcellular compartments where they perform diverse functions regarding cell cycle, metabolism and signal transduction ^11^. A number of these functions are carried out through intracellular calcium (Ca^2+^) fluxes ^12^. The endoplasmic reticulum (ER) is a welcoming hub for a number of Bcl-2 proteins where they interact with ER-localized Ca^2+^ channels and impinge on Ca^2+^ homeostasis regulation. For instance, Bcl-2, *via* its BH4 domain, interacts with the modulatory and transducing region (MTD) and ligand-binding region (LBR) of the inositol trisphosphate receptor 1 (IP3R1) thus inhibiting ER Ca^2+^ discharge ^13^ +and PMID: 30989245. In contrast, Bcl-xL interacts with the coupling domain (CD) of IP3R. Bcl-xL sensitizes the channel to low concentration of IP3. Hence, this interaction induces ER Ca^2+^ release and promotes Ca^2+^ oscillations required for mitochondrial energetics ^14^. Of note, mouse embryonic fibroblasts (MEFs) in which Bcl-xL is exclusively targeted to the ER were reported to be capable of restoring Ca^2+^ homeostasis, even though the underlying mechanisms are not elaborated ^15^. In fact, although Bcl-xL has been reported to contribute to major processes governing cancer progression ^4-10^ and embryonic development ^16^, the respective contributions of Bcl-xL subcellular pools in these processes remain unclear.

To address this matter, we generated genetically modified mice by Cre-LoxP recombination system in which Bcl-xL is targeted exclusively either to the ER or to the mitochondria. Both recombinant mice survive the embryonic stage, suggesting that the ER localization of the protein is indeed crucial for embryonic development. Upon cytotoxic drug treatment, MEFs with Bcl-xL targeted to the ER were found to be more prone to cell death inducers but, interestingly, resisted to ER stress inducers. We thus hypothesized that Bcl-xL might confer this resistance through its influence on IP3R-Ca^2+^ release. Indeed, in this study, we show that ER-targeted Bcl-xL decreases ER Ca^2+^ release through IP3R during ER stress. In an interesting manner, ER-addressed Bcl-xL was found to reduce the Unfolded Protein Response (UPR) markers’ expression in ER stressful conditions, compared to wild type (WT) and mitochondria-targeted Bcl-xL. Altogether, our data shed the light on a moonlighting function of Bcl-xL at the ER where it dampens the UPR through the reduction of IP3R-dependent Ca^2+^ release, acting thus as potent regulator of ER stress.

## Results

### Bcl-xL silencing sensitizes to inducers of the unfolded protein response

Bcl-xL is an anti-apoptotic Bcl-2 family member known to repress Bax dependent MOMP. Indeed, bclx-knockout (KO) MEFs were found more sensitive to the protein kinase C (PKC) inhibitor Staurosporine, an activator of the mitochondria-dependent apoptosis pathway, as shown by using Sytox Green™ and Caspase 3 assays (Figure 1A-D & Supplementary Figure 1A). In the course of this study, we also observed that these KO cells were more sensitive to the sarco/endoplasmic reticulum Ca^2+^ ATPase (SERCA) inhibitor Thapsigargin (Figure 1A-D & Supplementary Figure 1B), indicating that, in addition to mitochondria-dependent apoptosis, *bclx* silencing may have consequences regarding ER-dependent processes. Indeed, Thapsigargin is a potent ER stress activator that signals through the unfolded protein response (UPR) suggesting a role of Bcl-xL in the ER stress response. Accordingly, we investigated the influence of Bcl-xL on the UPR by analyzing the expression profile of typical UPR markers ^17^. In an interesting manner, eIF2α expression and eIF2α phosphorylation were increased, even in the absence of stress. Moreover, CHOP and ATF4 levels were gradually and significantly enhanced in KO MEFs upon Thapsigargin treatment (Figure 1E) supporting that the ER pool of Bcl-xL could be an inhibitor of the UPR, in particular following treatment with ER stress inducers.

**Figure 1.**
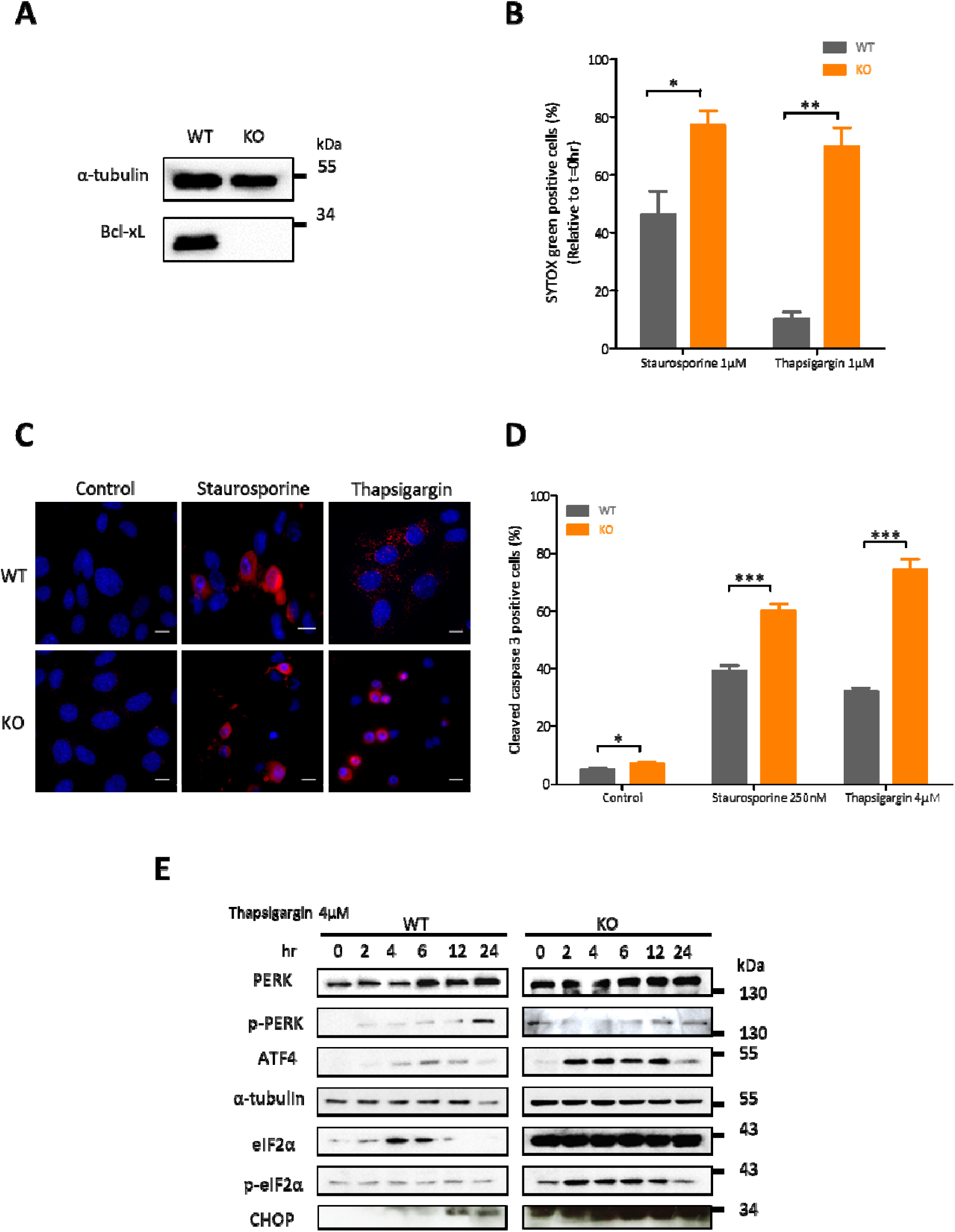
Bcl-xL protects from ER stress. (A) Western blot detecting Bcl-xL endogenous levels in WT and bclx KO MEFs. α-Tubulin was used as a loading control. (B) Cell death quantification (% of Sytox GREEN™marked cells) in WT and bclx KO MEFs treated with 1µM Staurosporine for 24hrs or 1µM Thapsigargin for 48 hrs (mean ± SEM; n=3; *, p<0.05; **, P<0.01). Results are displayed at t=15hrs and t=32hrs, respectively. C) Representative images of cleaved Caspase 3 stained WT and KO MEFS after Staurosporine and Thapsigargin treatment for 6hrs and 24hrs, respectively. Blue fluorescence: Nuclei. Red fluorescence: cleaved Caspase 3. Scale bar: 10µm. (D) Quantification of Cleaved Caspase 3 stained WT and KO MEFs after treatment with 250nM Staurosporine for 6hrs or 4µM Thapsigargin for 24hrs. 5 images of an average of 30 cells each were taken per cell type per experiment (n=3). Control: treated with DMSO (mean ± SEM; n=3; *, p<0.05; ***, p<0.001). (E) Kinetics of UPR markers in the presence of 4μM Thapsigargin for 24hrs in WT and *bclx* KO MEFs. α-Tubulin was used as a loading control.

### Generation of single subcellular localization mutants of Bcl-xL

The above data suggest that Bcl-xL is a negative regulator of ER stress, we thus analyzed its subcellular localization in MEFs isolated from C57BL/6 mice embryos. Subcellular fractionation experiments on WT MEFs highlighted the multiple subcellular localizations of Bcl-xL at the mitochondria, the cytosol and the ER (Supplementary Figure 2A).

**Figure 2.**
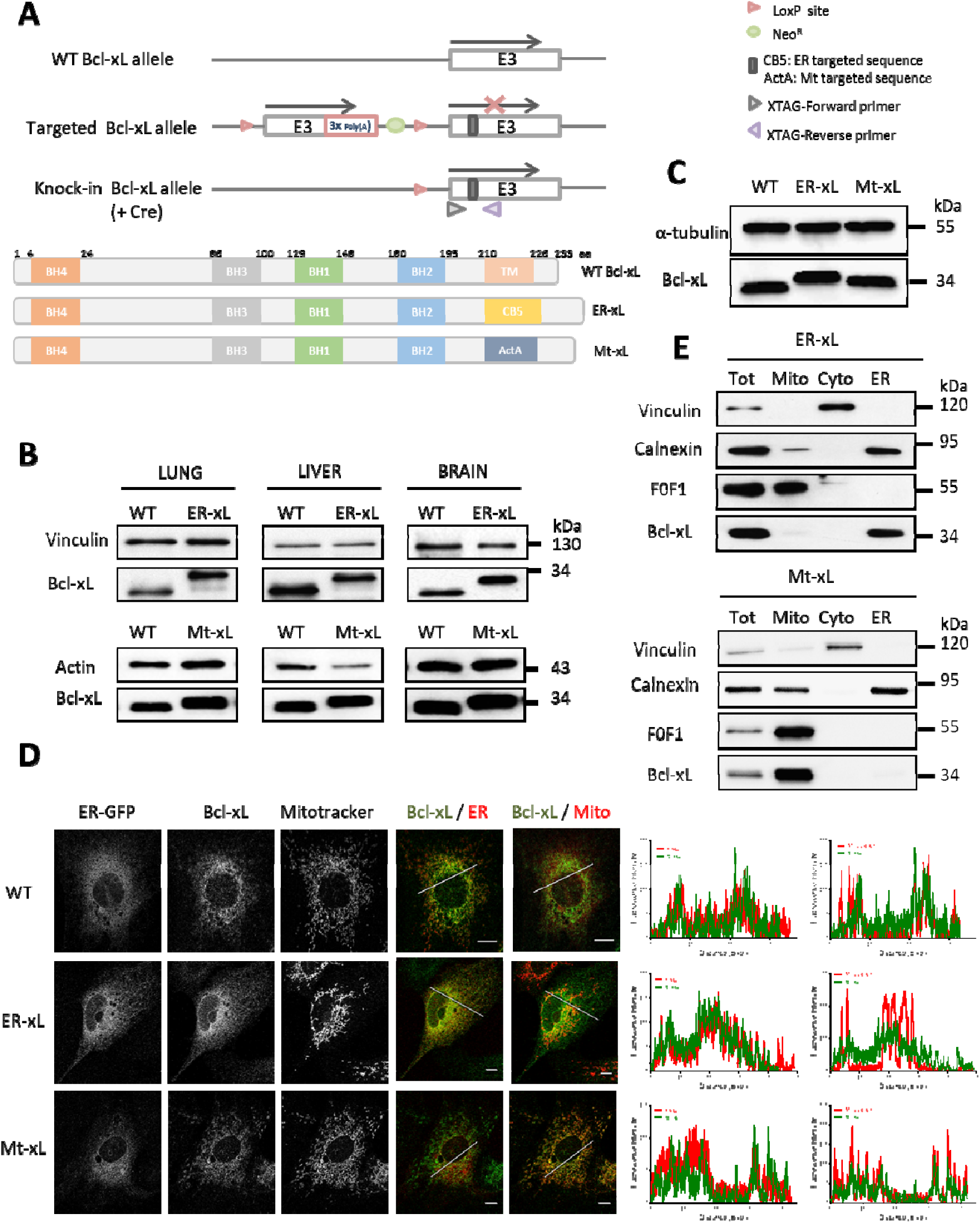
Generation and analysis of ER-xL and Mt-xL mice. (A) The mice genotyping strategy was done with a Cre-LoxP recombination system where an engineered vector containing the WT Bcl-xL exon 3 (E3) and CB5 or ActA mutant sequences targeting Bcl-xL either to the ER or mitochondria, respectively, was inserted to replace the WT Bcl-xL E3 in floxed mice. Cre-recombination induced WT exon 3 excision and CB5 or ActA mutant expression. Accordingly, three protein products are possible: WT Bcl-xL with intact TM domain, ER-xL with CB5 targeting sequence and Mt-xL with ActA targeting sequence. Neomycin is used as an inducer of vector resistance (Neo^R^). Two primers were used: XTAG-Forward and XTAG-Reverse. Poly(A): Poly(A) tail. (B) Western blot detecting Bcl-xL endogenous levels in cells extracted from the lung, liver and brain of WT, ER-xL and Mt-xL mice. Vinculin and actin were used as loading controls. (C) Western blot detecting Bcl-xL endogenous levels in WT, ER-xL and Mt-xL MEFs extracted at E13. α-tubulin is used as a loading control. (D) Bcl-xL subcellular localization detection in WT, ER-xL and Mt-xL MEFs by Immunofluorescence. Co-localization was assessed using ER-EGFP transfection and Mitotracker staining for ER and mitochondrial localization, respectively. Profile plots of fluorescence signals detecting the intensity of fluorescence along the white segments on merged images were quantified by ImageJ software and shown to the right. Scale bar: 10µm. (F) Bcl-xL endogenous expression in ER-xL (Top panel) and Mt-xL (lower panel) MEFs post-subcellular fractionation was detected by western blot. Vinculin is used as a cytosol marker, Calnexin as an ER marker and F0F1 ATPase as a mitochondrial maker. (Tot) whole-cell lysates; (Mito) mitochondria; (Cyto) cytosol.

In order to decipher the functional contribution of the ER and mitochondrial pools of Bcl-xL, we generated recombinant C57BL/6 mice by the Cre-LoxP system in which Bcl-xL is exclusively at the ER (ER-xL) or at the mitochondria (Mt-xL). An engineered vector containing the WT Bcl-xL exon 3 and Cytochrome B (CB5) or ActA mutant sequences that respectively target proteins to the ER or the mitochondria, was inserted to replace the WT Bcl-xL exon 3 in floxed mice. Cre-dependent recombination resulted in WT exon 3 excision and CB5 or ActA mutant expression. Accordingly, three protein products could be generated: WT Bcl-xL with intact transmembrane (TM) domain, ER-xL with CB5 targeting sequence and Mt-xL with ActA targeting sequence (Figure 2A).

Recombinant products were validated at both genomic and protein levels using polymerase chain reaction (PCR) and western blot, respectively (Supplementary Figure 2B). It should be noted that ER-xL (245 aa) and Mt-xL (242 aa) are slightly longer than WT Bcl-xL (233 aa). Bcl-xL expression was verified at the protein level in the lung, the liver and the brain of WT, ER-xL and Mt-xL mice (Figure 2B). Likewise, Bcl-xL protein expression was checked in MEFs derived from WT, ER-xL and Mt-xL embryos at embryonic day E13 (Figure 2C). These MEFs were immortalized by serial passaging and used in all experiments thereafter.

We next aimed at verifying the subcellular localization of Bcl-xL in the immortalized MEFs. To do so, we performed immunofluorescence experiments on all three types of MEFs transfected with ER-targeted EGFP. As shown in (Figure 2D), Bcl-xL is found at the ER in ER-xL MEFs, at the mitochondria in Mt-xL MEFs and co-localized in both organelles in WT MEFs, as indicated by the profile plots of fluorescence intensity. Subcellular localization of Bcl-xL was confirmed by fractionation (Figure 2E) and ImmunoGold labelling (Supplementary Figures 2C, D).

### Bcl-xL at the ER, and not at the mitochondria, confers resistance to ER stress

Upon verification of the expression and localization of Bcl-xL, we wanted to challenge the anti-apoptotic potential of the protein by treating the MEFs with cell death inducers. Accordingly, treatment for 6hrs with Staurosporine enhanced Caspase 3 activity in ER-xL (69.2±3.3% activated caspase 3-positive cells) MEFs compared to WT (43.1±2.7% activated caspase 3-positive cells) and Mt-xL MEFs (45.7±2.1% activated caspase 3-positive cells) see Figure 3A & Supplementary Figure 3A. These data were further confirmed by analyzing PARP cleavage (Figure 3B). Together these observations show that ER-xL MEFs exhibit higher susceptibility towards Staurosporine treatment. Of note, fluorescence activated cell sorting (FACS) measurements displayed higher subG1 population in ER-xL MEFs compared to Mt-xL and WT MEFs, confirming the cell death vulnerability of these cells (Supplementary Figure 3B).

**Figure 3.**
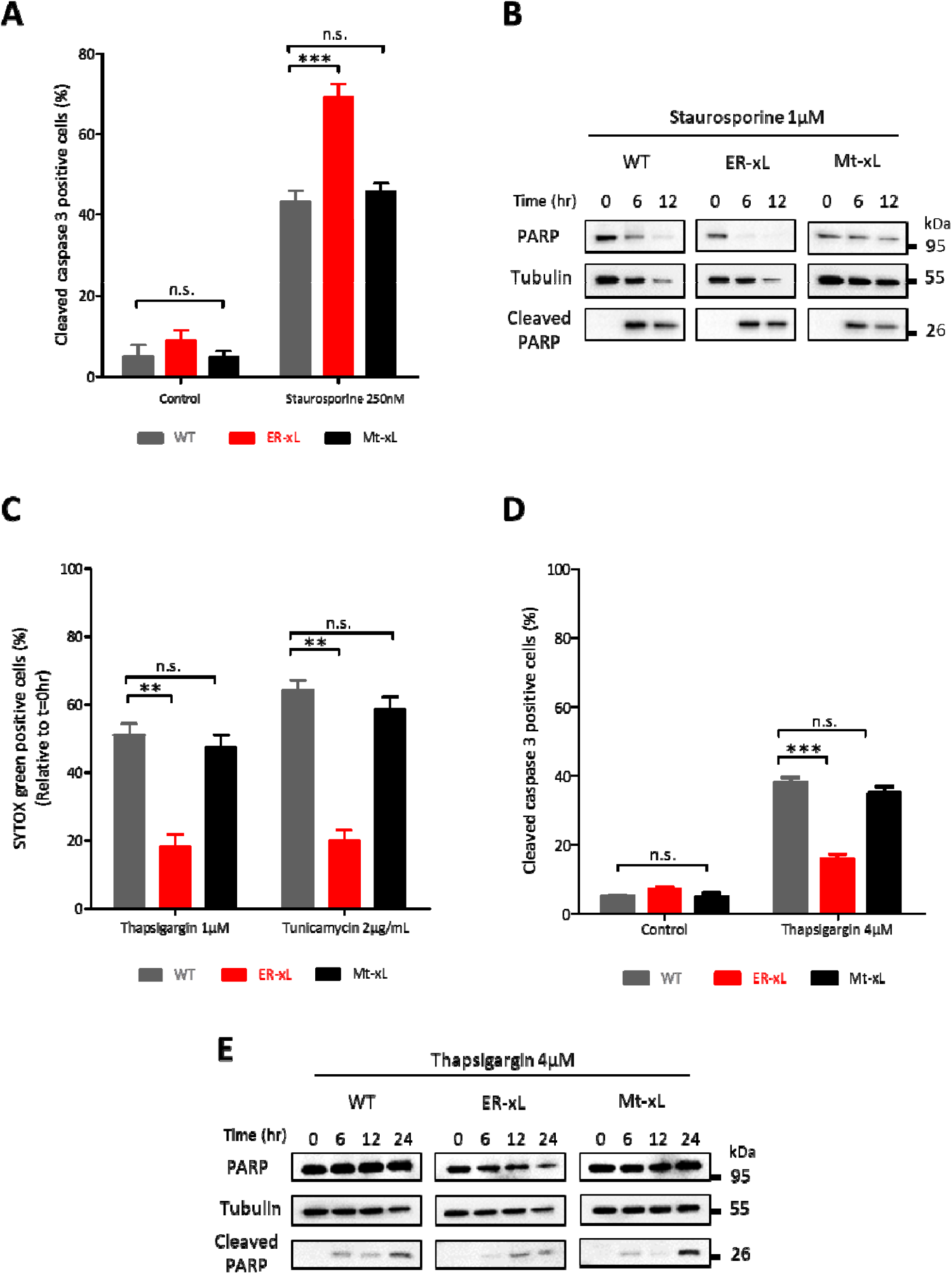
ER-xL MEFs resist ER stress-mediated cell death. (A) Quantification of cleaved Caspase 3 stained WT, ER-xL and Mt-xL MEFs after treatment with 250nM Staurosporine for 6 hrs. 5 images of an average of 30 cells each were taken per cell type per experiment (n=3). Control: treated with DMSO (mean ± SEM; n=3; ***, p<0.001; n.s., non-significant, p>0,05). (B) Kinetics of PARP cleavage in WT, ER-xL and Mt-xL MEFs following 1µM Staurosporine treatment over 12 hours. α-Tubulin was used as a loading control. (C) Cell death quantification (% of Sytox GREEN™-positive cells) in WT, ER-xL and Mt-xL MEFs treated with 1µM Thapsigargin or 2µg/mL Tunicamycin for 72 hrs (mean ± SEM; n=3; n.s., non-significant, p>0.05; **, P<0.01). Results are displayed at t=72hrs. (D) Quantification of cleaved Caspase 3 stained WT and KO MEFs after treatment with 4µM thapsigargin for 24hrs. 5 images of an average of 30 cells each were taken per cell type per experiment (n=3). Control: treated with DMSO (mean ± SEM; n=3; n.s., non-significant, p>0.05; ***, p<0.001). (E) Kinetics of PARP cleavage in WT, ER-xL and Mt-xL MEFs following 4µM Thapsigargin treatment over 24 hours. α-Tubulin was used as a loading control.

Next, we treated ER-xL and Mt-xL MEFs with the ER stress inducers Thapsigargin and Tunicamycin and evaluated cell death by measuring the percentage of Sytox Green™-positive cells as well as activated caspase 3-positive cells. Interestingly, ER-xL MEFs significantly resisted ER stress induction, compared to control and Mt-xL MEFs (Figure 3C,D & Supplementary Figures 3C-E). These observations were confirmed by measuring PARP cleavage (Figure 3E).

Altogether, these data indicate that ER-xL MEFs are more resistant to ER stress inducers than Mt-xL MEFs, whereas in contrast they are more prone to activate the mitochondria-dependent intrinsic apoptosis pathway.

### ER-localized Bcl-xL impairs UPR initiation under ER stress conditions

To further understand how ER-localized Bcl-xL protects from ER stress, we analyzed the expression of the UPR markers. Upon Thapsigargin treatment for 24hrs, ER-xL MEFs displayed reduced PERK phosphorylation compared to WT and Mt-xL cells (Figure 4A). This phosphorylation is in fact the first step of the PERK arm pathway activation in the UPR ^19^. Moreover, Mt-xL MEFs exhibited much higher ATF4 expression levels, and significantly increased phosphorylation of eIF2α and CHOP, compared to ER-xL MEFs and WT MEFs (Figure 4B).

**Figure 4.**
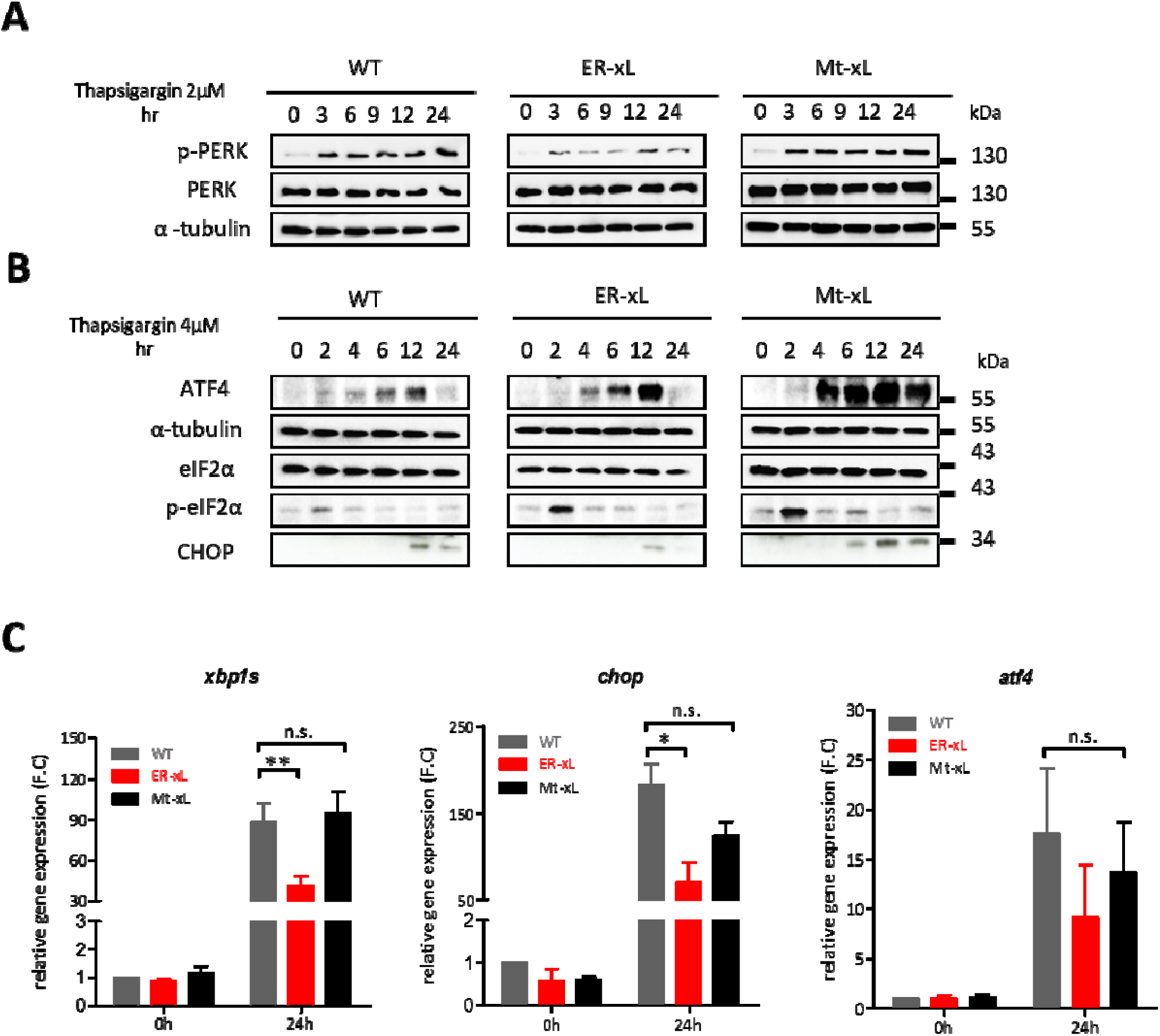
Bcl-xL at the ER reduces the UPR. Kinetics of UPR markers in the presence of 2μM (**A**) and 4μM (**B**) Thapsigargin over 24hrs in WT, ER-xL and Mt-xL MEFs. α-tubulin was used as a loading control. (**C**) RT-PCR assay was used to analyze the mRNA levels of the UPR markers *xbp1s, chop* and *atf4* in WT, ER-xL and Mt-xL MEFs after 4μM thapsigargin treatment over 24 hours.

Then, we ought to check whether the UPR response was also modified at the transcriptional level. In effect, during the UPR, ATF3, a downstream effector of ATF4, induces the transcription of CHOP and GADD34 ^20^. The latter is needed to guide protein phosphatase 1 (PP1) to dephosphorylate eIF2α. As for ATF5, it is a downstream effector of phosphorylated eIF2α that delays translation initiation upon ER stress ^21^. In addition, the IRE1 arm of the UPR results in *xbp1* splicing, thus activating the transcription of genes involved in cell death ^22^. Actually, ER-Bcl-xL, but not Mito-Bcl-xL, was found to decrease the mRNA levels of all above cited UPR actors (namely xbp1s, CHOP, ATF4), following Thapsigargin insult, in accordance with previously observed increased protein levels (Figure 4C, to be compared to Figure 1E). Collectively, these results support the notion that ER-based Bcl-x can potentially act as an inhibitor of the UPR stress response.

### ER-based Bcl-xL inhibits IP3R-Ca^2+^ release

In order to understand how Bcl-xL could act on the UPR pathway mechanistically, we hypothesized that this influence could be through ER Ca^2+^ regulation. Indeed, Bcl-xL is presumably a potent modulator of Ca^2+^ fluxes at the ER as it was previously reported to physically interacts with the IP3R coupling domain (CD) through its hydrophobic pocket. Moreover, Bcl-xL was recently shown to perform a biphasic regulation of IP3R gating at the ER, whereby low [Bcl-XL] sensitized IP3Rs while high [Bcl-XL] inhibited IP3Rs ^14,23,24^.

First, we demonstrated that Bcl-xL-IP3R interaction actually existed in WT MEFs. Indeed, Proximity Ligation Assay (PLA) using antibodies targeting specifically Bcl-xL and IP3R detected significantly higher number of PLA dots in WT MEFs (18.6±1.7%) compared to KO cells (0.80±0.2%) (Figure 5A,B).

**Figure 5.**
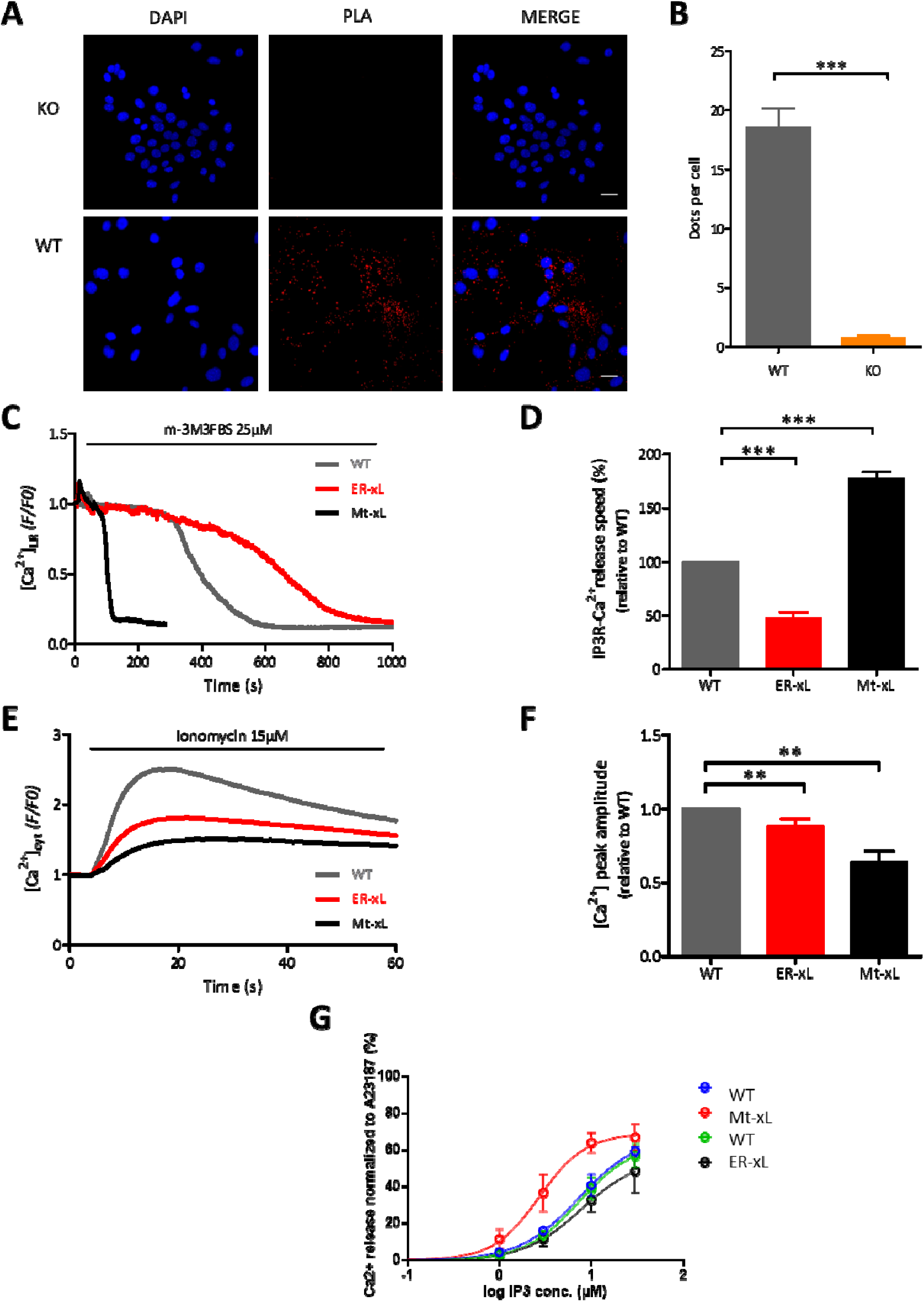
Bcl-xL at the ER interacts with IP3R and inhibits its activity. (A) Representative images showing the endogenous interactions between IP3R - C terminus and Bcl-xL in WT and KO MEFs after Proximity ligation assay (PLA). DAPI was used to stain nuclei. Scale bar: 30µm. (B) Quantification of PLA experiments (mean ± SEM; n=3; ***, p< 0.001). (C) Representative curves of ER calcium release in WT, ER-xL and Mt-xL MEFs transfected with CEPIA-1er after 25μM m-3M3FBS (PLC activator) injection. (D) Quantification of the slope coefficient of ER calcium release after PLC activation in MEFs (mean ± SEM n=3; ***, p< 0.001). (E) Representative curves of cytosolic calcium release assessed with 5μM Fluoforte after 15μM Ionomycin injection in WT, ER-xL and Mt-xL MEFs using a plate reader. (D) The ratio of fluorescence indicating [Ca^2+^] peak amplitude relative to WT is shown (mean ± SEM; n=3; **, p<0.01). (G) Representative curves of ER calcium release normalized to A23187 maximal ionophore release response after ^45^Ca^2+^ flux analysis. IP3-induced calcium release in Mt-Bcl-xL MEFs is clearly sensitized compared to wild-type MEFs and ER-Bcl-xL.

Furthermore, we evaluated the effect of Bcl-xL on passive ER-Ca^2+^ discharge in ER-xL and Mt-xL MEFs. To this end, ER-Ca^2+^ passive leakage (in the absence of IP3 stimulation) was analyzed following Thapsigargin treatment in MEFs transfected with R-CEPIA1er. As shown, ER-xL MEFs showed higher Ca^2+^ leakage (120.9±1.9%) compared to WT control MEFs (100%), contrary to Mt-xL MEFs (101.7±1.5%), see Supplementary Figure 4A,B. Interestingly, this increase was coupled with the activation of store operated Ca^2+^ entry (SOCE), an important mechanism that promotes the entry of Ca^2+^ from the extracellular space to the ER in response to ER-Ca^2+^ depletion ^27,28^. In ER-xL MEFs, SOCE was markedly increased (5.9±0.3%) relative to WT MEFs (3.9±0.3%). Of note, this process was also enhanced in Mt-xL MEFs (5±0.4%), see Supplementary Figure 4C,D.

We then evaluated the capacity of ER-xL to control IP3 dependent Ca^2+^ release. To do so, we activated IP3R by enhancing the production of IP3 through the Phospholipase C activator m-3M3FBS and measured the decrease of the fluorescent ER-targeted Ca^2+^ indicator R-CEPIA1er fluorescence. IP3R-dependent Ca^2+^ release in ER-xL MEFs was significantly slower (46.8±6.1%) relative to WT control MEFs (100%), see Figure 5C,D. This inhibition of Ca^2+^ release from the ER led to concomitant decrease in cytosolic Ca^2+^ levels in ER-xL (80±5%) relative to WT control MEFs (100%), see Figure 5E,F. Of note, Mt-xL MEFs exhibited higher rates of IP3R-dependent Ca^2+^ release (177.5±6%), see Figure 5D, together with limited cytosolic Ca^2+^ increase (70±6%) compared to control (Figure 5F), which is consistent with previous studies about the role of mitochondrial Bcl-xL on VDAC-dependent Ca^2+^ uptake ^23^.

To directly assess the impact of Bcl-xL on IP3R-mediated calcium release, we performed ^45^Ca^2+^ flux assays on plasma membrane-permeabilized MEF cells expressing WT Bcl-xL, ER-Bcl-xL or mt-Bcl-XL. In these experiments, ER Ca^2+^ stores are loaded to steady state with ^45^Ca^2+^, after which unidirectional passive Ca^2+^ leak from the ER is initiated by thapsigargin. This is quantified as fractional loss (%/2min) as a function of time. Once a constant fractional loss is obtained, cells are challenged with varying concentrations of IP3. The total releasable Ca^2+^ is determined by using A23187, a Ca^2+^ ionophore. IP3-induced Ca^2+^ release, normalized to A23187, was strongly sensitized for cells expressing mt-Bcl-xL compared to WT-Bcl-xL, while it was similar for ER-Bcl-xL compared to WT-Bcl-xL (Figure 5G). These data indicate that Bcl-XL present at ER membranes, either in WT Bcl-xL or ER-Bcl-xL-expressing cells, inhibits IP3Rs in comparison to Bcl-xL present at mitochondrial membranes. In this experimental analysis, IP3-induced Ca^2+^ release was similar for WT Bcl-xL versus ER-Bcl-xL, which may indicate that the majority of WT Bcl-xL is already present at ER membranes and inhibits IP3Rs. As such, mt-Bcl-xL rather serves as an “Bcl-xL-knockout at the ER”.

### ER-xL decreases the UPR by inhibiting IP3R-mediated Ca^2+^ depletion upon ER stress induction

Together our results highlight, first, the protection conferred by ER-Bcl-xL against ER stress and second, the inhibition by ER-Bcl-xL of IP3R-dependent ER Ca^2+^ release. To confirm that the effect of Bcl-xL on Ca^2+^ fluxes is actually IP3R-dependent, we inhibited the IP3R by 2-Aminoethoxydiphenyl-borate (2-APB) ^29,30^ and analysed ER Ca^2+^ release in MEFs upon Thapsigargin treatment for 24hrs. Actually, a significant reduction in UPR markers’ expression was observed in both WT and Mt-xL MEFs upon treatment with 2-APB (Figure 6A). The use of Xestospongin C, another IP3R inhibitor ^31,32^, confirmed these observations (Figure 6B). Thus, although these inhibitors are not strictly specific for IP3R (2-APB also targets TRP Ca2+ channel of the TRP family ^33,34^, potassium channels ^35^, and volume-regulated anion channels^36^, whereas as Xestospongin C also inhibits SERCA ^37,38^), collectively our observations suggest that the UPR is at least in part dependent on IP3R Ca^2+^ permeability.

**Figure 6.**
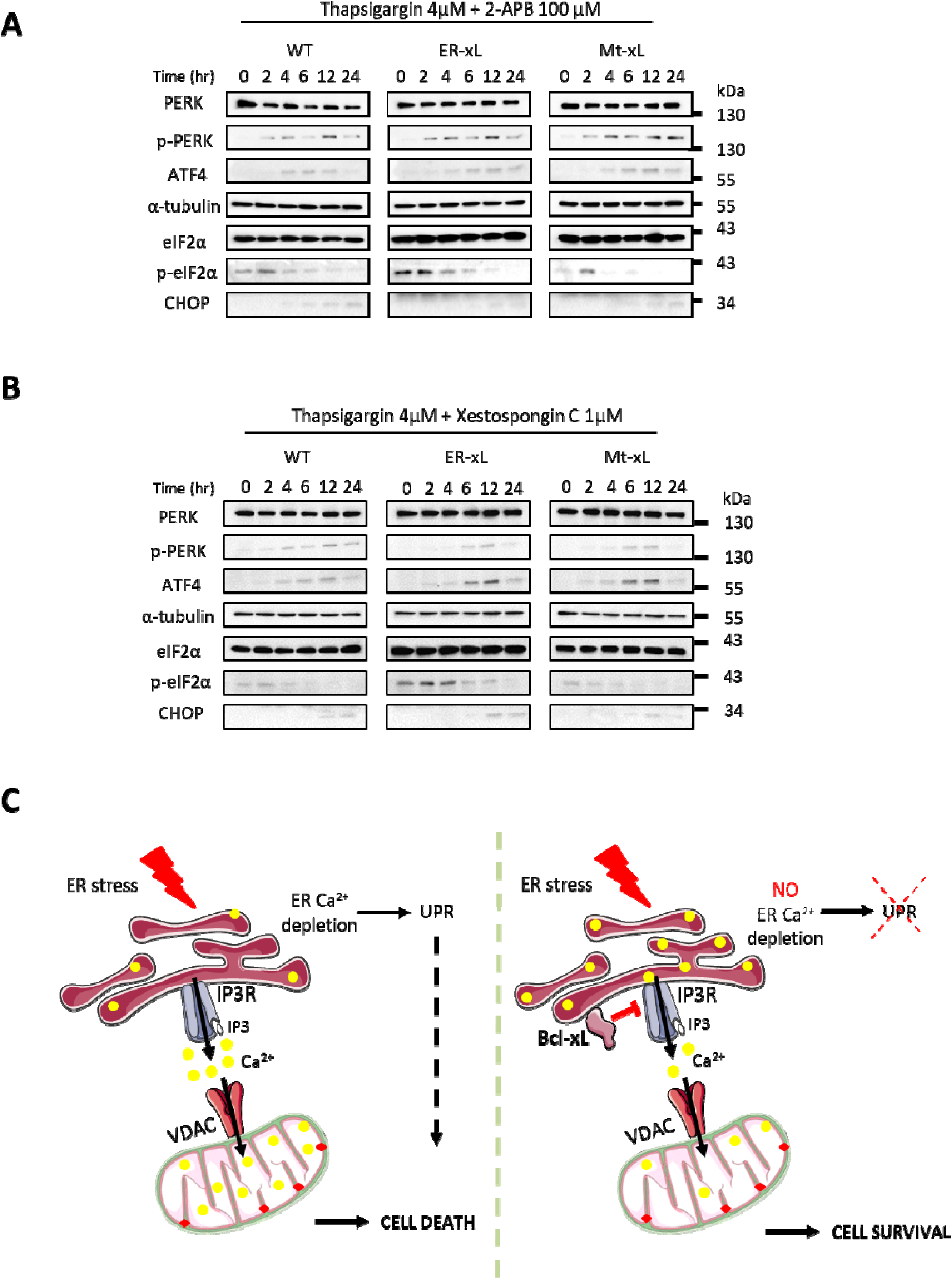
Bcl-xL at the ER reduces the UPR by inhibiting IP3R upon ER stress induction. (A) Kinetics of UPR markers after 4µM Thapsigargin and 100µM 2-APB treatment over 24 hours in WT, ER-xL and Mt-xL MEFs. 2-APB is considered as an IP3R inhibitor. α-Tubulin was used as a loading control. (B) Kinetics of UPR markers after 4µM Thapsigargin and 1µM Xestospongin C treatment over 24 hours in WT, ER-xL and Mt-xL MEFs. Xestospongin C is an IP3R and SERCA inhibitor. α-Tubulin was used as a loading control. (C) Proposed model for the role of Bcl-xL at the ER. ER stress might deplete ER Ca^2+^ stocks resulting in: 1-mitochondrial apoptosis initiation through mPTP opening and 2-the UPR activation leading to the transcription of genes promoting apoptosis. Hence, the outcome is cell death. Under these conditions, Bcl-xL at the ER interacts with and inhibits the IP3R abrogating ER Ca^2+^ depletion required for apoptosis induction. Consequently, Bcl-xL at the ER performs a new indirect anti-apoptotic function upon ER stress.

## Discussion

In addition to their role in regulating the mitochondrial pathway of apoptosis, non-canonical functions of Bcl-2 proteins emerged throughout the years ^12^. Some of these functions are in part ensured through intracellular Ca^2+^ fluxes regulation ^37,39^. Indeed, although mitochondria are considered the major hub of Bcl-2 proteins, a number of Bcl-2 homologs, including Bcl-2 itself, can also localize to the ER where they exert opposing effects on ER-mediated Ca^2+^ handling ^40^. Regarding Bcl-xL, although mainly involved in the mitochondrial pathway of apoptosis ^41^, a number of reports support that it contributes to a range of other processes, independent of apoptosis, including autophagy, migration, metabolism and signal transduction ^40,42^. Some of these non-canonical roles of Bcl-xL are presumably performed at the level of the ER ^14,43^ although such a subcellular localization has long been controversial. Indeed, Bcl-xL TM domain was reported to address Bcl-xL at the mitochondria, but not at the ER ^41^, wherease in contrast, Bcl-xL was shown to interact with the ER-based IP3R1 Ca^2+^ channel.

As an anti-apoptotic protein, Bcl-xL is a promising target regarding cancer therapy ^7,8,10^. In this respect, understanding the molecular mechanisms that drive its subcellular localization may help prevent unwanted side effects of Bcl-xL inhibitors, including BH3 mimetics ^44,45^. Here, we specifically interrogated mechanisms beyond the localization of Bcl-xL at the ER and their involvement in ER Ca^2+^ turnover. bclx KO MEFs’ vulnerability to cytotoxic drugs replicated previous data about MOMP prevention by Bcl-xL ^2^. Furthermore, we observed that bclx KO enhanced the UPR, following Ca^2+^-dependent insults, suggesting that Bcl-xL might be involved in the control of ER stress.

By generating knock in mice expressing either ER- or mitochondria-targeted Bcl-xL, we aimed at further understanding the respective roles of both Bcl-xL subcellular pools. Staurosporine significantly enhanced cell death in ER-xL MEFs, compared to Mt-xL MEFs, further confirming the cell death protection potential of mitochondrial Bcl-xL ^15^. Interestingly, ER-xL MEFs were found to be more resistant to the ER stress inducing agents Thapsigargin and Tunicamycin, compared to Mt-xL MEFs, which was corroborated by reduced UPR markers expression. Of note, Bcl-xL was previously shown to protect cells from ER stress by sequestering Bim at the mitochondria, and subsequently reducing CHOP-induced apoptosis during plasma cell differentiation ^46^. In addition, mitochondrial Bcl-xL was shown to inhibit the translocation of Bim to the ER upon ER stress thereby abrogating caspase 12-mediated apoptosis in murine myoblast cells ^47^. Here, we show that ER-addressed Bcl-xL contributes also to ER stress silencing.

At the molecular level, Bcl-xL was reported to interact with the C-terminal domain of IP3R in the neighborhood of its channel pore. Through this interaction, Bcl-XL sensitizes IP3Rs to basal IP3, thereby promotingER-to-mitochondria Ca^2+^ signaling ^14,22^. At higher Bcl-XL concentrations, Bcl-XL could also bind to the coupling domain of IP3Rs, thereby provoking inhibition of Ca2+ flux through the channel (22). We confirmed these observations using PLA in MEFs. Moreover, our Ca^2+^ fluxes measurements showed that ER-xL actually promotes passive ER Ca^2+^ release in non-stressful conditions, which presumably maintains mitochondrial Ca^2+^ uptake and mitochondrial bioenergetics ^48^. In contrast, Ca^2+^ fluxes measurement performed in the presence of high IP3 levels supported the notion that ER-xL prevents the full opening of IP3R Ca^2+^ channels in stressful conditions. This phenomenon might act as a survival defense mechanism in case of ER stress by limiting ER Ca^2+^ stores depletion and downstream UPR initiation. Actually, based on the duration and severity of the stress, unfolded proteins accumulate in ER lumen with concomitant decrease in ER Ca^2+^ levels. As a consequence, ER-stress sensors (PERK, IRE1 and ATF6) are activated, setting in motion adaptive responses that prompt either cell survival or cell death, depending on the severity and the duration of the stress ^49^.

Considering a link between the inhibitory action of Bcl-xL on IP3R and the resistance towards ER stress observed in ER-xL MEFs, we hypothesized that the direct closure of IP3R, using the dedicated pharmacological agents 2-APB and Xestospongin C, should also reduce UPR markers expression in WT and Mt-xL MEFs, by limiting luminal Ca^2+^ decrease which was indeed found to be the case.

Finally, we propose a novel mode of action of the anti-apoptotic member Bcl-xL at the ER (Figure 6 C). According to this model, upon ER stress induction, Bcl-xL would negatively regulate IP3R-mediated Ca^2+^ release and prevent ER Ca^2+^ depletion. Thus, in this way Bcl-xL would avert UPR initiation and downstream apoptosis, further promoting cell survival.

## Materials and methods

### Cell lines and drugs

Mouse Embryonic Fibroblasts (WT, ER-xL and Mt-xL) were extracted from mice embryos at Embryonic day 13 (E13) and cultured along with bclx KO MEFs (kindly provided from Li laboratory ^15^) under standard cell culture conditions (37°C, 5% CO) in Dulbecco’s modified Eagle’s medium (DMEM) high glucose medium (Gibco) supplemented with 10% Fetal Bovine Serum (FBS ; Sigma Aldrich), 100U/mL penicillin and 100μg/mL streptomycin (Gibco). Several cytotoxic drugs, Ca^2+^ ionophores, channel inhibitors were used: Staurosporine (Sigma Aldrich), Etoposide (Sigma-Aldrich), Thapsigargin (Enzo Life Sciences), Tunicamycin (Sigma Aldrich), Ionomycin (Enzo Life Sciences), 2-APB (Tocris), Xestospongin C (Tocris), m-3M3FBS (Sigma Aldrich).

### Cell death detection

For cell death detection experiments, MEFs were analyzed using Incucyte ZOOM live-cell imaging system (Essen Bioscience). In brief, cell death was assessed using SYTOX Green™ nucleic acid stain at 250nM concentration (ThermoFisher Scientific) plus the addition of several cell death inducers: Staurosporine 1μM, Etoposide 10μM, Thapsigargin 1μM, Tunicamycin 2μg/mL. Images were automatically acquired in phase and green fluorescent channels every one hour for a maximum of 72 hours (hrs) at 4X magnification. Data were processed using a dedicated algorithm (Essen Bioscience). Cell proliferation was estimated using the phase-contrast images of the controls of the same experiments.

### Immunofluorescence

MEFs were seeded on a glass coverslip in 12-well plates at around 40% of confluence and transfected with ER-targeted EGFP. After cell attachment, the medium was discarded and the cells were incubated with mitochondria-staining dye (MitoTracker Red CMXRos, Life technologies) for 20min at 37°C then fixed with 4% Paraformaldehyde with 0.3% Triton X100. MEFs were washed 3 times with 0.1% Triton-PBS then incubated for 20 minutes (min) with blocking buffer (0.1% Triton X100, 3% BSA in PBS). Cells were subsequently incubated for 1hr with primary antibodies and 1hr with Alexa fluor 563 coupled secondary antibodies. Cells were washed 3 times after the incubation with both antibodies. Coverslips were mounted with a Dako mounting medium. Bcl-xL antibody (Cell Signaling #2764; 1/200) was used to detect endogenous Bcl-xL. For apoptosis assay after 250nM Staurosporine treatment for 6hrs or 4µM Thapsigargin treatment for 24hrs, the same procedure was performed without transfecting the MEFs. Cleaved Caspase 3 (Cell Signaling #9661, 1/1000) antibody was used to assess cell death followed by Alexa fluor 563 coupled secondary antibodies. Nuclei were detected with Hoechst 33342 dye (Invitrogen #H3570) at 10μg/mL concentration. Images were acquired using a Zeiss 780 confocal microscope.

### Antibodies used for Western Blot

The following antibodies were used: Vinculin (Santa Cruz #sc-55465; 1/2000), Purified Mouse Anti-ATP Synthase β – F0F1 (BD Transduction Laboratories #612518; 1/2000), Calnexin (Cell signaling #2679; 1/1000), α-Tubulin (Santa Cruz #sc-32293; 1/1000), Actin (Sigma-Aldrich #A2066; 1/500), PERK (Cell signaling #3192; 1/1000), phospho-PERK (Cell signaling #3179; 1/1000), eIF2α (Cell Signaling #2103, 1/1000), phospho-eIF2α (Cell Signaling #3398, 1/1000), ATF4 (Cell Signaling #11815, 1/1000), CHOP (Cell Signaling #5554, 1/1000), Bcl-xL (Cell Signaling #2764, 1/1000), PARP (abcam #ab6079, 1/500), Cleaved Caspase 3 (Cell Signaling #9661, 1/1000). HRP-conjugated goat anti-mouse and goat anti-rabbit antibodies (DAKO) were used as secondary antibodies. Immunoblotting was performed according to standard procedures. For kinetic immunoblotting, MEFs were treated with the indicated drug and were harvested at each indicated time point where they were lysed and their protein content was measured.

### Subcellular fractionation

MEFs were harvested by trypsinisation and homogenized in MB buffer (210mM mannitol, 70mM sucrose, 1mM EDTA and 10mM HEPES (pH 7.4) containing protease inhibitors) by shearing (around 20 times) with a 1mL syringe and a 26G 5/8 needle. Homogenization was visually confirmed under the microscope. All steps were carried out at 4°C. Cell extracts were centrifuged 3 times (10min at 1,500g) to remove the nuclear fraction (pellet). Supernatants were then centrifuged 3 times (10min at 10,600 g). After the third round of centrifugation, crude mitochondria pellet was carefully collected with a micropipette, suspended in MB buffer and a Percoll medium (225mM mannitol, 25mMHEPES (phH 7.4), 1mMEGTA and 30% Percol (vol/vol)) and centrifuged for 30min at 95000g separating the MAMs and the mitochondria. The supernatant was centrifuged for 1hr at 100,000g. After this last round of centrifugation, the obtained microsome pellet was resuspended in RIPA buffer. The purity of obtained fractions was assessed by Western blot.

### Intracellular calcium measurements

For ER-Ca^2+^ measurements, MEFs were transfected with 2µg RCEPIA 1er probe 48hrs before starting the experiment. Cells were seeded at 40% confluence in 8 well Nunc™ Lab-Tek™ II Chamber Slide and incubated with EM buffer (121mM NaCl; 5.4mM KCl, 2.6mM MgCl2 hexahydrate, 6mM NaHCO3, 5mM D-Glucose, 25mM HEPES pH 7.3). 10µM Thapsigargin injections were done while quantifying the fluorescence with a Zeiss LSM 780 confocal microscope.

SOCE measurements were performed on MEFs incubated for 1hr with 5µM Fluoforte dye, and washed with BBS containing 0.1mM EGTA. After 20 seconds (s) of baseline fluorescence reading, Thapsigargin 10µM was injected followed by 2mM CaCl_2_ once the fluorescence drops back to baseline

For cytosolic Ca^2+^ measurements, MEFs were seeded onto a 96-well plate one day before the experiment. Cells were then loaded with 5µM Fluoforte dye (Enzo Life Sciences) in EM buffer for 45min at 37°C followed by 2 washes with EM buffer. Fluorescence was assessed using the ClarioSTAR plate reader, following 15µM Ionomycin injections (Enzo Life Sciences)

For IP3R-induced Ca^2+^ release, transfected MEFs with RCEPIA 1er probe were injected with 25 µM m-3M3FBS (Sigma-Aldrich), direct activator of PLC. The Fluorescence was assessed using Zeiss LSM 780 confocal microscope. The speed of Ca^2+^ release was calculated by measuring the slope of the increase of fluorescence upon injection.

### ImmunoGold labelling

Subcellular localization of endogenous Bcl-xL in WT, ER-xL, Mt-xL and bclx KO MEFs was performed by Bcl-xL ImmunoGold labeling and Transmission Electron Microscopy (TEM) imaging. Uranyle acetate staining was used to allow for subcellular structure visualization in fixed cell sections. Images were collected using a JEOL 1400JEM transmission electron microscope and Zeiss LSM780 confocal microscope.

### Proximity Ligation Assay (PLA)

MEFs were grown on coverslips and fixed with methanol for 2 min. After two PBS washes, MEFs were saturated with the blocking solution followed by a 1hr at 37°C incubation with the primary antibodies: Bcl-xL (Cell Signaling #2764, 1/1000) and IP3R C-terminal (abcam #ab190239, 1/1000). PLA secondary antibodies probes were added and incubated for 1hr at 37°C after 3 PBS washes. Ligation and amplification (100 min at 37°C) steps followed. Cells were mounted with Duolink II Mounting medium containing DAPI and images were acquired using the fluorescence microscope.

### ^45^Ca^2+^ assay

^45^Ca^2+^ flux analysis was performed as previously described^25,26^. After permeabilizing the MEFs, IP3R-mediated Ca^2+^ release was assessed through the addition of increasing concentrations of IP3. Steady state ^45^Ca^2+^ were used to load intracellular non-mitochondrial Ca^2+^ stores where the release of ^45^Ca^2+^ from the plasma membrane was assessed at specific time intervals. 1.5µM maximal IP3 concentration was used. MEFs were loaded with ^45^Ca^2+^ in the presence of the Ca^2+^ ionophore A23187 in order to determine the passively bound Ca^2+^ value that will be subtracted to calculate exclusively the amount of releasable Ca^2+^ from the IP3R.

### Cell cycle FACS analysis

FACS analysis with Propidium Iodide (PI) (abcam) was carried out by flow cytometry (BD FACSCanto™ II, BD Bioscience) according to manufacturer’s recommended protocol. Briefly, MEFs were seeded and grown overnight onto a 12-well plate. Afterwards, they were trypsinized and harvested. The obtained cell pellet was washed with PBS and fixed with cold ethanol for 30min at 4°C. After 2 PBS washes, MEFs were treated with 50 µl of a 100 µg/ml stock of RNase to ensure that only DNA is stained. 200 µL of PI (from 50 µg/ml stock solution) was then added and the cell cycle was analyzed by flow cytometry (BD FACSCanto™ II, BD Bioscience) using untreated cells as negative control for gating. A total of 10000 events were recorded during the experiments (n=3). The percentage of each cell population was analyzed using BD FACSDIVA 8.0.1 software (BD Bioscience, USA).

### Genotyping by PCR

PCR-based analysis was used to genotype mice. Each reaction mixture consisted of deoxynucleoside triphosphates, MgCl_2_, a forward primer specific for Bcl-xL exon 3: EXE-F (5’-GGAAAGGCCAGGAGCGCCTTC-3’), a reverse complementary primer specific for Bcl-xL exon 3: XTAG-R (5’-CCCAACCCTGTGATAGGGCAAG-3’), Taq polymerase, specific buffer and genomic DNA. Conditions for PCR were as follow: (1) 5min at 98°C; (2) 35 cycles, with 1 cycle consisting of 30s at 94°C, 30s at 59°C, and 40s at 72°C and (3) an elongation period of 5min at 72°C. Amplified products were analyzed by electrophoresis in 2% agarose gels and bands were visualized by Gel-Doc BIO-RAD imaging system.

### RT qPCR

Total RNA was isolated from MEFs following treatment with 4µM thapsiragin over 24hrs with the NucleoSpin RNA plus from MACHEREY-NAGEL, following the manufacturer’s instructions. For the reverse transcription, 1µg was used for cDNA synthesis using a Moloney Murine Leukemia Virus (MMLV) reverse transcriptase (SuperScript II from ThermoFisher Scientific) and random primers. One mL of RT reaction was used for the PCR following the protocol: 1 min 94uC and 20 cycles of amplification by 1 min 94uC, 1 min 55uC, 1 min 72uC, and finishing with 5 min at 72uC. Amplified products were visualized in 1% agarose with ethidium bromide. The following primers were used:

### Statistical analyses

Student t-tests were performed when comparing two means and ANOVA was used for comparison of three means or more. A p-value equal to or under 0.05 was considered statistically significant.

## Acknowledgements

We would like to acknowledge Stéphane Borel for skillfull technical assistance. We are thankful for their help to staff members from Anican (Centre Léon Berard) and Alecs (SFR Lyon Est) mouse facilities as well as Cicle cell imaging facility (SFR Lyon Est). This work was supported by grants from AFM (# 20269, to GG) and Comité du Rhône de la Ligue contre le Cancer (to NP). LJ is a fellow from Ministère de l’enseignement supérieur et de la recherche. TN is a fellow from Inserm (Plan Cancer Initiative)

