## Supplementary figures S1-4 for "The Endoplasmic Reticulum pool of Bcl-xL dampens the Unfolded Protein Response through IP3R-dependent Calcium Release"

**Supplementary Figure 1**

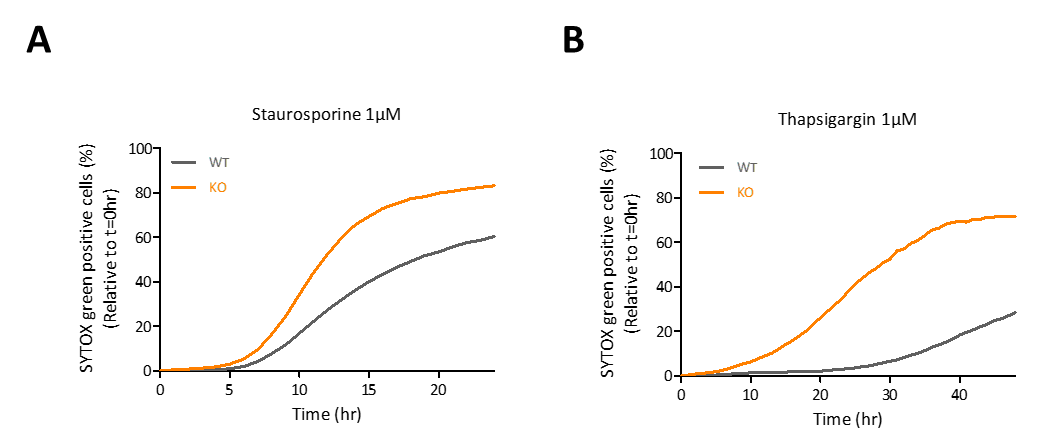

**Supplementary Figure 1. Cytotoxic and ER-stress mediated drug treatment on WT and KO MEFs**

Representative curves of cell death quantification (% of Sytox GREEN^TM^ marked cells) in WT and *bclx* KO MEFs treated with Staurosporine 1µM (A) or or with thapsigargin 1µM (B) for 48 hrs. SYTOX Green^TM^ is used to detect dead cells.

**Supplementary Figure 2**

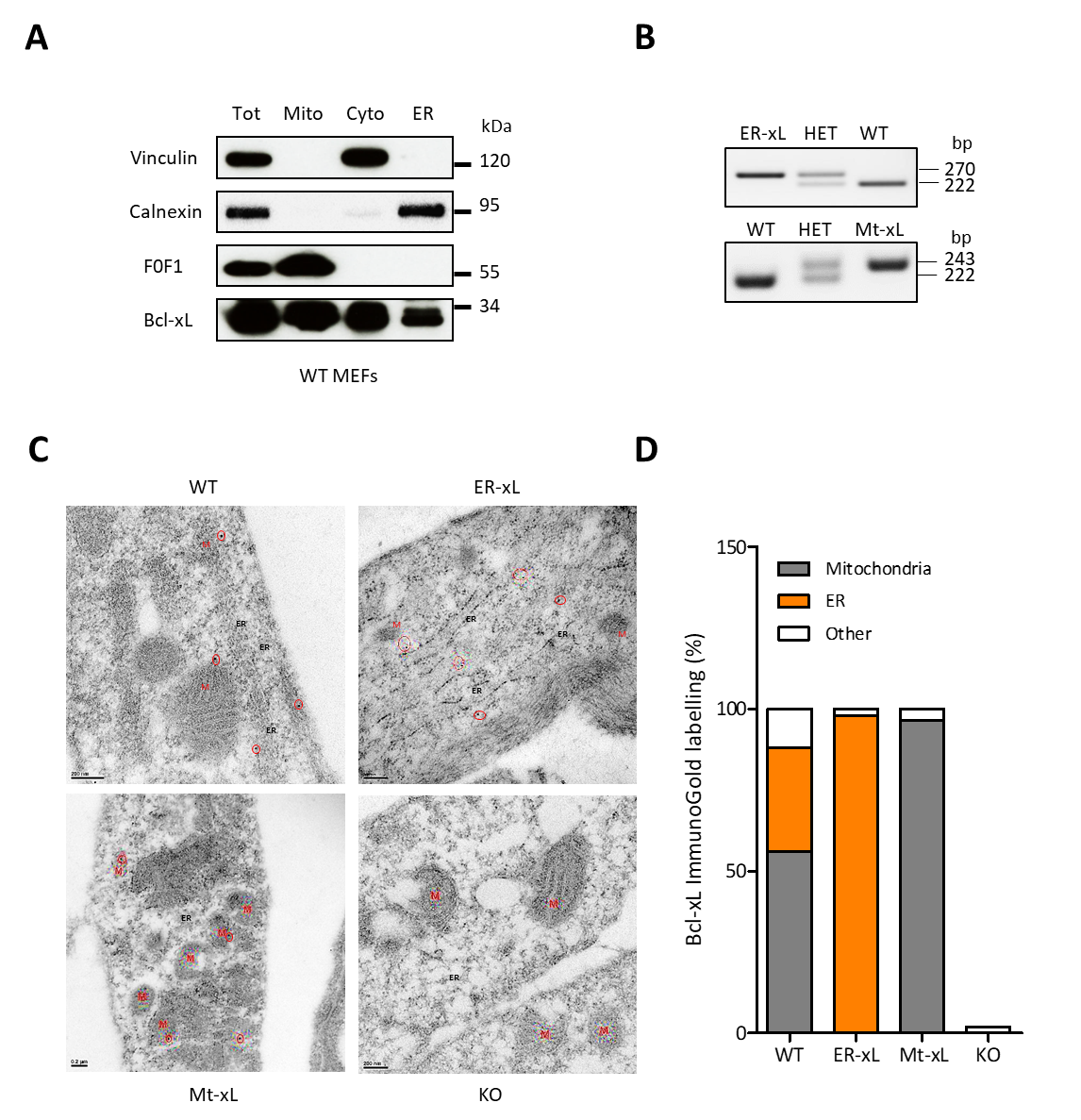

**Supplementary Figure 2. Genotyping and localization of Bcl-xL**

(A) The localization of endogenous Bcl-xL in WT MEFs was assessed by subcellular fractionation resulting in four fractions: (Tot) whole-cell lysates; (Mito) mitochondria; (Cyto) cytosol and ER followed by western blot where vinculin is used as a cytosol marker, Calnexin as an ER marker and F0F1 ATPase as a mitochondrial maker. (B) PCR results confirming the genotype of the genetically modified mice: WT for Bcl-xL^(WT/WT)^; HET for Bcl-xL^(WT/ActA)^ or ^(WT/CB5)^; ER-xL for Bcl-xL^(CB5/CB5)^ and Mt-xL for Bcl-xL(^ActA/ActA)^. Bp: base pair. HET: Heterozygous. (C) Representative images for Bcl-xL ImmunoGold labeling in WT, ER-wL, Mt-xL and *bclx* KO MEFs. Red circle: Gold particle; M: mitochondria; ER: Endoplasmic Reticulum. Scale bar: 200nm. (D) Quantification of Bcl-xL ImmunoGold labelling in WT, ER-wL, Mt-xL and *bclx* KO MEFs (n=15 cells per cell type).

**Supplementary Figure 3**

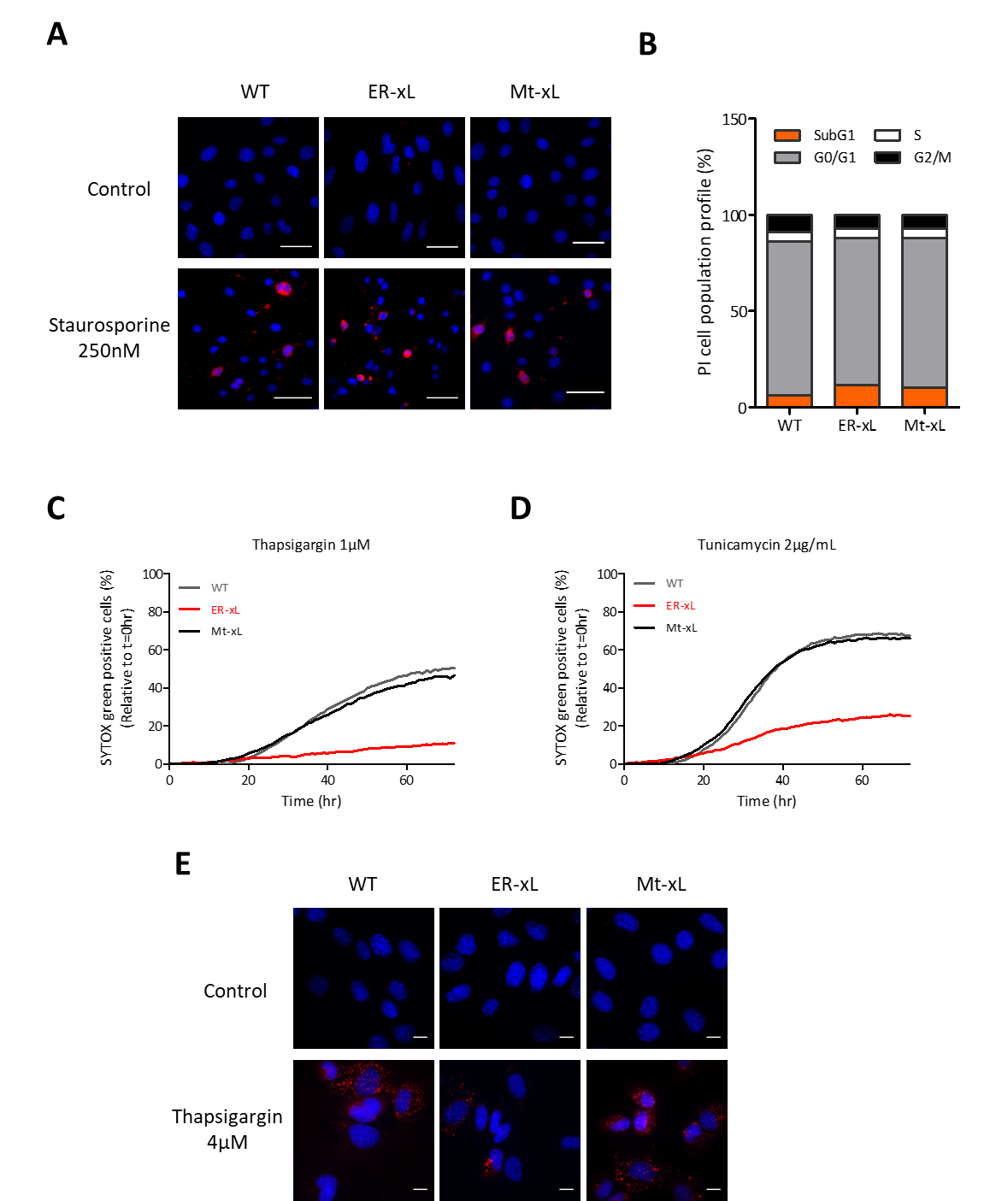

**Supplementary Figure 3. Bcl-xL at the ER protects from ER stress induced cell death**

(A) Representative images of cleaved Caspase 3 stained WT, ER-xL and Mt-xL MEFS after staurosporine treatment for 6hrs. Blue fluorescence: Nuclei. Red fluorescence: cleaved caspase 3. Scale bar: 50µm (B) Percentages of DNA content in each cell cycle phase are shown for each cell type after FACS analysis with Propidium Iodide (PI) in WT, ER-xL and Mt-xL MEFs (n=3). Representative curves of cell death quantification (% of Sytox GREEN^TM^-positive cells) in WT, ER-xL and Mt-xL MEFs treated with Thapsigargin 1µM (C) and Tunicamycin 2µg/mL for 72 hrs (D). SYTOX Green^TM^ is used to detect dead cells. (E) Representative images of cleaved caspase 3 stained WT, ER-xL and Mt-xL MEFS after Thapsigargin treatment for 24hrs. Blue fluorescence: Nuclei. Red fluorescence: cleaved caspase 3. Scale bar: 10µM.

**Supplementary Figure 4**

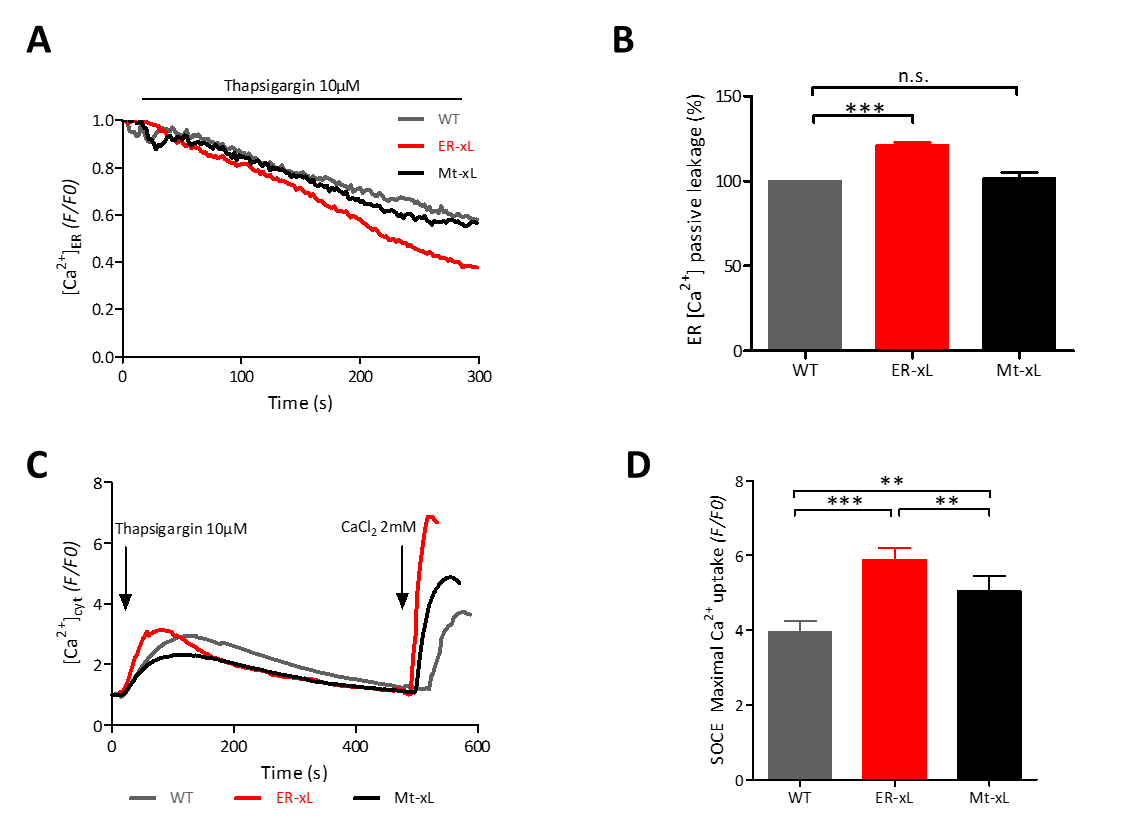

**Supplementary Figure 4. Bcl-xL at the ER affects intracellular Ca^2+^ fluxes.**

(A) Representative curves of ER passive calcium leakage in WT, ER-xL and Mt-xL MEFs transfected with CEPIA-1er after 10μM Thapsigargin injection. (B) Quantification of the slope coefficient of ER passive calcium leakage in MEFs (mean ± SEM; n=3; ***, p< 0.001; n.s., non-significant). (C) Representative curves of Store Operated Calcium Entry (SOCE) assessed with 5 μM Fluoforte after draining the ER from calcium by 10μM Thapsigargin injection followed by 2mM CaCl_2_ injection in WT, ER-xL and Mt-xL MEFs. (D) The ratio of fluorescence indicating the maximal calcium uptake after CaCl_2_ injection is shown (mean ± SEM; n=3; **, p<0.01; ***, p< 0.001).
